# Targeted pre-conditioning and cell transplantation in the murine lower respiratory tract

**DOI:** 10.1101/2024.11.12.622518

**Authors:** Anat Kohn, Michael J. Herriges, Payel Basak, Liang Ma, Bibek R. Thapa, Darrell N. Kotton, Finn J. Hawkins

## Abstract

Transplantation of airway basal stem cells could achieve a durable cure for genetic diseases of the airway, such as cystic fibrosis and primary ciliary dyskinesia. Recent work demonstrated the potential of primary- and pluripotent stem cell (PSC)-derived basal cells to efficiently engraft into the mouse trachea after injury. However, there are many hurdles to overcome in translating these approaches to humans including developing safe and efficient methods for delivery in larger animal models. We propose a model which targets preconditioning and cell-delivery to the intrapulmonary airways utilizing a micro- bronchoscope for delivery. The detergent polidocanol was adapted for distal lung pre-conditioning, inducing intrapulmonary airway epithelial denudation by 5 and 24-hours post-delivery. While initial re- epithelialization of airways occurred later than tracheas, complete repair was observed within 7-days. Both PSC-derived and primary basal cells delivered via micro-bronchoscope post-polidocanol injury engrafted in tracheas and intrapulmonary airways, respectively. Transplanted cells differentiated into ciliated and secretory lineages while maintaining a population of basal cells. These findings demonstrate the utility of bronchoscopically targeted pre-conditioning and cell delivery to the conducting intra- pulmonary airways, providing an important framework for pre-clinical translation of approaches for engineered airway epithelial regeneration.

## Introduc4on

Cystic Fibrosis (CF) and primary ciliary dyskinesia (PCD) are monogenic diseases which result in significant morbidity and mortality. Although they arise from distinct cellular defects, both CF and PCD result in chronic lung disease characterized by impaired mucociliary clearance, recurrent respiratory infections, bronchiectasis, and ultimately end-stage lung disease. While highly effective modulator therapies have significantly improved the lives of most persons with CF, there are currently no targeted therapies for a subset of CF patients with Class I mutations or any patients with PCD. Apart from supportive therapies, lung transplantation is the only therapy available to these individuals when they develop end-stage lung disease. However, availability of donor lungs is limited, and the procedure is associated with significant morbidity and poor long-term survival. In adults, 1- and 5-year survival post- lung transplantation are 85.3% and 54.3% respectively; by contrast, 1- and 5-year survival post-heart transplantation are 90.8% and 80% respectively (1, 2). Cell-based therapy represents a promising theoretical alternative for airway epithelial defects such as CF and PCD, in which a patient’s epithelial cells would be replaced through engraftment of genetically normal primary or induced pluripotent stem cell (iPSC)-derived cells into their own lungs. Importantly, through the use of gene-edited autologous tissue-resident stem cell transplantation, this treatment strategy would circumvent the need for immunosuppression while providing a durable cure of the underlying disease though these efforts are in the very early stages of development (3, 4).

In the airways, basal cells are the major tissue resident stem cell, having both self-renewal and mul0-lineage differentiation potential (5–7). Further, both mouse and human primary or iPSC-derived basal cells can be expanded in culture and differentiated into other airway lineages (6, 8–14). These traits make basal cells the leading candidate for durable airway cell transplantation as they can generate all the specialized cell types of the airway epithelium while maintaining the stem cell pool (15). In a recent study our group demonstrated the engraftment of both primary and pluripotent stem cell (PSC)-derived basal cells (15) into syngeneic mouse tracheas. Engraftment was dependent on pre-conditioning the tracheas to remove the majority of the endogenous epithelial cells so that donor cells can attach, proliferate, and then differentiate. This was achieved through the delivery of the detergent polidocanol (PDOC) via oropharyngeal aspiration (OPA). (15–17). Following transplantation, donor-derived-cells persisted for over two years and were able to differentiate into ciliated and secretory populations while maintaining a basal stem cell pool. Importantly, these donor-derived stem cells maintained their stem cell function, as demonstrated by their ability to proliferate and differentiate in response to subsequent rounds of injury as well as repopulate a secondary donor trachea in serial transplantation studies for at least eight generations to date (15). This maintenance of stemness is critical in developing a durable cell- based transplantation therapy. Finally, in proof-of-concept experiments, these same techniques were utilized to engraft primary human bronchial epithelial cells (HBECs) and human induced pluripotent stem cell (iPSCs) derived basal cells (Hu-iBCs) into the tracheas of NOD scid gamma (NSG) immunodeficient mice (15). Interestingly, despite the cross-species differences, the human basal cells persisted in the tracheas of NSG mice for at least 6 weeks and differentiated into secretary and ciliated lineages.

These studies demonstrate the potential of airway epithelial cell-based therapies; however, clinical translation of this approach faces many hurdles that include the: 1) development of preconditioning methods that are effective but safe, 2) demonstration that donor cells can actually reverse a disease phenotype, and 3) demonstration of safety and efficacy in larger animal models. To this end, we hypothesized that targeted preconditioning and cell-delivery will be required for larger animal models and sought to develop this first in the mouse polidocanol model of engraftment. The field of cell-based therapies for the lung is in its infancy. The majority of intrapulmonary epithelial cell transplantation has focused on alveolar engraftment, with the only other reported airway engraftment seen following injury with Naphthalene as an injury model (18–23). While these studies demonstrate the potential of engrafting cells in the airways, a systemically delivered conditioning agent with known toxicity is not ideal moving into large animals and humans. An intrabronchial conditioning/injury agent delivery by bronchoscopy is a logical approach to targeting specific lung lobes or segments and to minimize lung damage and systemic effects. The use of bronchoscopy to target a single lobe for therapeutic injury is already in use in humans in the seong of thermal ablation for severe persistent asthma(24, 25). However, accessing the intrapulmonary airways in mice is a challenge, due to their small size (mouse trachea inner diameter 1.5-2.5 mm) (26, 27). Recently, a micro-bronchoscope was described by Dames et al, who demonstrated targeted delivery of fluids, infectious particles (bacteria and viruses), as well as allergens into the murine IPAs. Schelde et al have similarly utilized a micro-bronchoscope to deliver bacteria embedded in agarose particles to the distal IPAs in a directed, unilateral manner to mimic chronic infections (28).

Here we propose a model of intrapulmonary airway pre-conditioning utilizing a micro- bronchoscope to deliver polidocanol in a targeted, unilateral manner. Initial epithelial repair has a slower onset than in the trachea following polidocanol injury. Basal cells delivered via micro-bronchoscope, after polidocanol injury, engraft in both the trachea and intrapulmonary airways, and differentiate into secretory and ciliated cells, while maintaining a population of basal cells. Together this demonstrates the utility of the micro-bronchoscope for precise epithelial injury and cell transplantation, providing an important framework for using existing bronchoscopy techniques to perform similar procedures in larger animal models and ultimately patients.

## Materials and Methods

### Animal maintenance

All mouse studies were approved by the Institutional Animal Care and Use Committee of Boston University Chobanian and Avedisian School of Medicine. All mice were purchased from Jackson Laboratory (Strains C56BL/6J #000664, UBC-GFP #004353, 129X1/SvJ 000691, and 129S1/SvImJ # 002448) and maintained in facilities overseen by the Animal Science Center at Boston University. Experiments were performed in 8-12 weeks old male and female mice.

### Cell culture of murine iBCs and primary BCs

We have previously detailed the culture, passaging, maintenance, and lentiviral tagging of murine PSC-derived basal cells (iBCs) in basal cell medium (BCM)(15, 29). In brief, murine PSC-derived basal cells (iBCs) were differentiated from Nkx2-1^mCherry^ mouse ESCs and labeled with a pHAGE-EF1aL-GFP-W (30) lentivirus as previously described (15, 31) and utilized from cryopreserved stocks, referred to throughout as “iBCs”. Primary BCs were collected and cultured from adult UBC-GFP mouse (Jackson Labs, Strain #004353) tracheas as previously described (15). Primary BC and iBCs were cultured in 3D Matrigel in BCM and cell digest was performed as previously described (15, 29).

### Fluorescence-ac4vated cell sor4ng (FACS) or analysis

Single-cell suspensions were prepared as previously described (15, 29). When necessary, cells were stained with NGFR antibody (Abcam, ab245134) for 30 minutes in FACS buffer (2% FBS in PBS) followed by incubation in secondary antibody for 20 min at 4C and then resuspended in FACS buffer with 1:100 DRAQ7 or 1:1000 Calcein Blue (live/dead stain). FACS was performed on a Beckman Coulter MoFLo Astrios and flow analysis was performed on a either a Beckman Coulter MoFLo Astrios or a Stratedigm S100TIEXi. Resulting plots were further analyzed using FlowJo v10.8.1.

### Micro-Bronchoscope

The micro-bronchoscope system (Polydiagnost, Germany) was previously described by Dames et al. It comprises modular endoscope, built-in monitor, xenon light source, camera (semi-rigid 3000-pixel optics, outer diameter 0.4mm), a modular bronchoscope handle with three proximal Luer connections and a distal Luer connection for the cannula (PolyShaft, outer diameter 0.75mm, length 5timm). The camera connects though the central proximal Luer and passes though the handle and through the cannula.

Fluids can be flushed though the lumen via one of the two side-port Luer connections (Supp Fig 1A). Assembly and use were performed based on the manufacturer’s instructions. For disinfection the system was initially flushed with Peroxigard (0.5% hydrogen peroxide disinfectant solution, Fisher #NC1739226), followed by 70% ethanol solution, then tissue-culture grade sterile water (Corning, 25055CV) to remove any residue of the disinfecting solutions, finally the working channel was dried by flushing with medical-grade air (Linde, AI M-AE).

**Figure 1.**
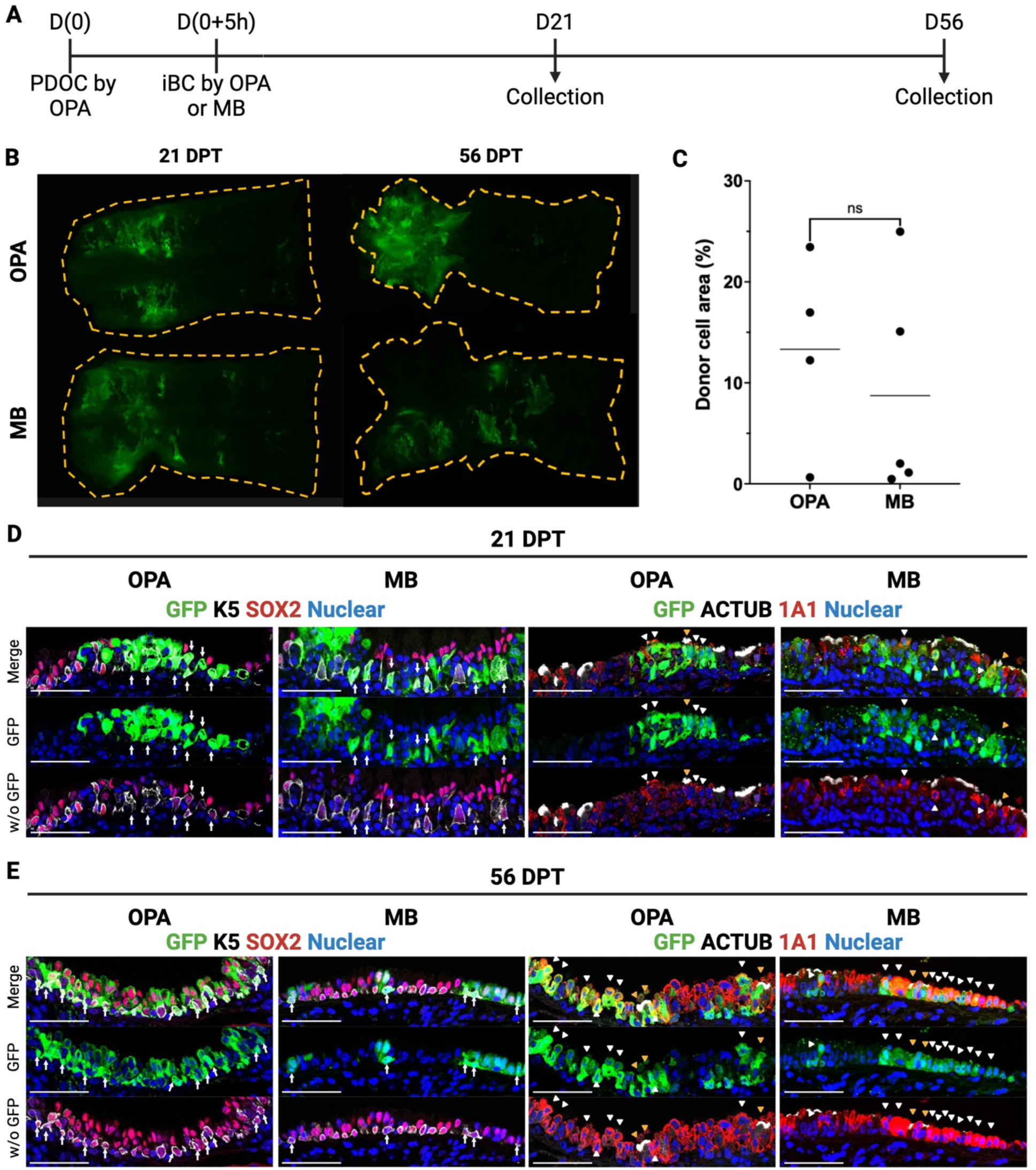
: EngraJment of cells delivered through a micro-bronchoscope (A) Schematic for transplantation of iBCs into mouse tracheas using oropharyngeal (OPA) or micro- bronchoscopic (MB) delivery methods following OPA delivery of polidocanol (PDOC). (B) Representative images of flayed tracheas 21 or 56 days post transplant (DPT). Trachea tissue is indicated by a dashed yellow line. (C) Quantification of the percent of the trachea surface area that is GFP+ following transplantation. n.s. = not significant by t-test. n=4,5 biological replicates. (D) Representative immunofluorescence confocal microscopy of trachea showing donor-derived cell clusters at 21 days post-transplantation, assessing colocalization of GFP with airway lineage markers including K5 (basal cells, white arrows), ACTUB (ciliated cells, yellow arrowheads), and SCGB1A1 (secretory cells, white arrowheads). Scale bars are 50 um. (E) Representative immunofluorescence confocal microscopy of trachea showing donor-derived cell clusters at 56 days post-transplantation, assessing colocalization of GFP with airway lineage markers including K5 (basal cells, white arrows), ACTUB (ciliated cells, yellow arrowheads), and SCGB1A1 (secretory cells, white arrowheads). Scale bars are 50 um.

For bronchoscopy, mice were anesthetized with ketamine (100 mg/kg BW; Covetrus, NDC 11695- 0703-1) and xylazine (50 mg/kg BW; Akorn, NDC 59399-110-20) via intraperitoneal injection (IP), then suspended by the front teeth by a wire attached to an angled plexi-glass support with a heating pad (HotHands® Hand Warmers) under the tail to maintain body temperature throughout the procedure. The tongue was gently retracted using fenestrated forceps (Roboz, RS-5160) and the micro-bronchoscope was introduced into the oral cavity and directed to the posterior oropharynx. After visualization of the vocal cords (Supp Fig 1B2) the micro-bronchoscope was advanced into the trachea (Supp Fig 1B3) and directed into the left or right mainstem bronchus. Post-procedure, mice were transferred into heated cages and monitored until recovered.

For application of fluids, one of the proximal Luer connection side port was capped (Cole-Parmer, 4550456) and the desired volume of fluid (20-100 ul) was pipetted into the remaining Luer connection side port followed by a slow flush of 250 cc air from a 1ml syringe. Fluid delivery was confirmed by visualization.

### Tracheal delivery of polidocanol and iBCs

For transplantation targeting the trachea, 2% polidocanol and iBCs were delivered by oral-pharyngeal aspiration as previously described (15, 16). In short, mice (males, 8-12 weeks old, progeny of a cross of 129X1/SvJ females with 129S1/SvImJ males) (Jackson Labs Strain #000691 and #002448, respectively) were anesthetized with 3% isoflurane until respiration slowed to 1 breath every 2-3 seconds, and 20 ul of 2% polidocanol was delivered directly into the posterior oropharynx (detailed in Ma et al. 2024).

Transplantation was conducted at 5 hours post-injury with 1.5-2 million congenic iBCs (described above) suspended in 30 ul PBS. Half of the injured mice received iBCs by oral-pharyngeal aspiration (29), the other through the working channel of the micro-bronchoscope. For transplantation targeting the trachea, the micro-bronchoscope was positioned in the proximal one third of the trachea just distal to the vocal cords.

### Mouse tracheal harvest and tracheal whole mount fluorescence imaging

Recipient mice were sacrificed via cervical dislocation following isoflurane overdose. Subsequent to PBS perfusion through the right ventricle mouse tracheas were harvested from larynx to carina and opened by longitudinal transection of the anterior trachea through the cartilaginous rings. For whole mount imaging, the transected trachea were placed on a glass slide and imaged by fluorescence microscopy using a Nikon Eclipse Ni-E microscope. Epifluorescence imaging quantification was performed with Image J (Version 2.1.0/1.53i). Specifically, the ‘‘Color Threshold’’ function was used to select and calculate the area of GFP+ regions (based on GFP signal) as well as the entire recipient trachea (based on background autofluorescence). Transplantation efficiency was calculated by dividing the GFP+ area over total trachea area. To prepare 7 mm thick tissue sections for immunostaining, the tracheal tissue was fixed with 4% PFA at 4C for 4 hours prior to processing and embedding in paraffin.

### Intrapulmonary airway polidocanol injury and primary basal cell transplanta4on

For PDOC delivery into the intrapulmonary airway, the tip of the micro-bronchoscope was placed within the left mainstem bronchus and 30 ul PDOC (0.25-1.75% diluted in PBS) was instilled as described above. Intrapulmonary polidocanol injury was characterized in both male and female C57BL/6J mice with similar response to injury.

For UBC-GFP primary basal cell transplants all recipients were C57BL/6J mice (Jackson Labs Strain #000664), both male and female with primary cells being isolated from the appropriate gender donor mice (Jackson Labs Strain #004353). Intrapulmonary PDOC (0.5%, 30 ul) was delivered to the left mainstem bronchus 24 hours prior to transplantation. On the day of transplantation donor cells were digested down to a single-cell suspension, as described above. These cells were suspended in Dual SMAD-Inhibitor media (DSI) and left in a 37C incubator for 2-3 hours, with flicking the conical every 30 minutes, to recover from digestion. At the end of this period cells were counted on a hemocytometer and resuspended in PBS (1.0-1.5e6 cells in 5tiul per mouse). These cell suspensions were delivered via micro-bronchoscopy to the left mainstem bronchus as described.

### Mouse lung harvest and lung whole mount fluorescence imaging

Mice were euthanized as described above and lungs were inflation-fixed with 4% paraformaldehyde (PFA) after PBS perfusion through the right ventricle. Both lungs and tracheas were removed en bloc and fixed overnight in 4% PFA. Whole mount fluorescent imaging of fixed lungs was conducted using a Nikon SMZ18 Stereomicroscope on freshly fixed samples. All samples were then processed, embedded in paraffin and sectioned at 8um. Hematoxylin and Eosin (H&E) stain was performed by standard protocols utilizing alcoholic Eosin Y. For immunohistochemistry, slides were blocked using normal donkey serum.

The M.O.M. blocking kit (Vector Laboratories, BMK-2202) was utilized when primary mouse antibodies were required. Sections were incubated with up to three primary antibodies (Supp Table 2) overnight at 4C. The next day slides were stained with Hoechst nuclear stain and up to three secondary antibodies (Supp Table 2) for 1 hour at room temperature and mounted with ProLong Diamond Antifade Mountant (Fisher, P36965). Stained slides were imaged using a Leica SP5 Confocal Microscope, Nikon Eclipse Ni-E microscope, or Akoya Biosciences Vectra Polaris Whole Slide Imaging System.

### Sta4s4cal methods

Relevant statistical methods are outlined in the figure legends. Unpaired, two-tailed Student’s t tests were used for comparisons involving only two groups, while ANOVA was used when considering multiple groups. GraphPad Prism 10 software was used to run statistical analysis.

## Results

### A micro-bronchoscope can be u4lized for murine airway visualiza4on and unilateral fluid delivery

We sought to recapitulate the key findings initially reported by Dames et al using the micro- bronchoscope (Sup Fig 1A) in mice. After induction of anesthesia and positioning of the mouse the micro-bronchoscope was advanced past the vocal cords (Supp Video 1, Supp Fig 1B2) and maneuvered into the trachea. Once at the main carina (Supp Fig 1B3) the micro-bronchoscope could be advanced into the left mainstem bronchi or into the right mainstem bronchi (Supp Video 1). In the right mainstem bronchus the bifurcation to the cranial lobe was easily identified (Supp Video 1) and the bronchoscope could be advanced further into the bronchus intermedius where the trifurcation to the three lower lobes were visualized (Supp Video 1, Supp Fig 1B4). From the bronchus intermedius 50 ul of green histologic dye was delivered through the working channel of the micro-bronchoscope under direct visualization. To confirm localization, paraffin sections were obtained, and counter stained with hematoxylin and eosin (H&E) (Supp Fig 1C). These confirmed the presence of green dye within the right-sided lobes (Supp Fig 1, C2-5) without evidence of dye in the left lobe (Supp Fig1, C1).

### Delivery of iBCs through the micro-bronchoscope does not alter engraJment ability in an established tracheal injury model

To ensure that airway basal cells could be delivered through the micro-bronchoscope working channel without significant loss of engraftment potential we utilized our established tracheal transplantation model of mouse pluripotent stem cell derived BCs (iBCs) post-polidocanol injury (15). Mouse iBCs containing an NKX2-1^mCherry^ reporter were labeled with a lentivirus constitutively expressing GFP. These cells were subsequently sorted to purify labeled iBCs based on co-expression of GFP, Nkx2-1^mCherry^, and NGFR. These cells were expanded for 2 weeks in BCM as airway organoids in Matrigel droplets prior to transplantation (Supp Fig 2A). On the day of transplant the percentage of GFP+;Nkx2-1^mCherry+^;NGFR+ cells was determined by flow cytometry (Supp Fig 2B,C). For transplantation, tracheas were first preconditioned through oropharyngeal aspiration (OPA) of 2% PDOC (15, 16), and then at 5-hours post-injury mice received donor cells (1.5-2e6 cells per mouse suspended in 30 ul PBS) delivered either via OPA or micro-bronchoscope (MB) targeted to the upper trachea (Fig 1A).

**Figure 2.**
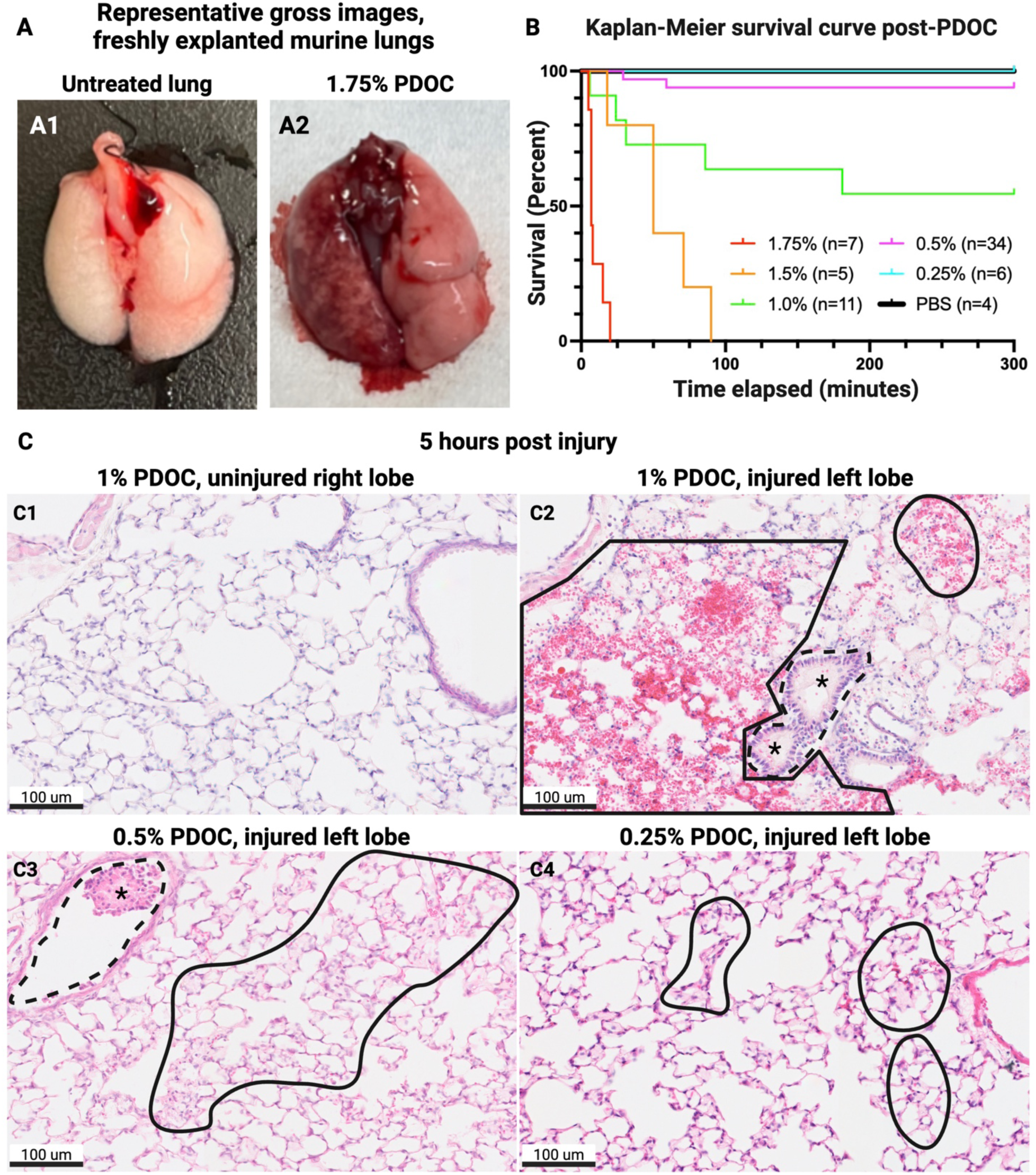
: Alveolar damage and hemorrhaging following high doses of polidocanol (A) Representative images of untreated lungs and hemorrhagic lungs following delivery of PDOC through a micro-bronchoscope. (B) Kaplan-Meier survival curve demonstrating survival following delivery of 3tiul of different percentage PDOC solutions. n indicates the number of mice per condition. (C) Representative H&E images of lungs 5 hours post-delivery of indicated percentage PDOC solutions highlighting airways (dashed lines), sloughed epithelial cells (asterisk), and alveolar damage (solid lines).

Mice were sacrificed at 21- and 56-days post transplantation (DPT) for analysis. Explanted tracheas were dissected along the cranio-caudal axis and visualized immediately post-harvest under wide field epifluorescence microscopy. At both time points GFP+ donor-derived cells could be readily observed whether delivered by OPA or MB (Fig 1B). When transplant efficiency was calculated by GFP+ area over total recipient tracheal area there was no statistical difference between the two groups (Fig 1C; OPA N=4, MB N=5). Similar to the OPA-delivered cells, MB-delivered cells maintained expression of airway epithelial markers NKX2-1 (data not shown) and SOX2 (Fig 1D, E), with a subset maintaining expression of the basal cell marker K5 at both 21 and 56 DPT (Fig 1 D,E white arrows). Furthermore, both MB- and OPA-delivered donor cells were able to differentiate into secretory and ciliated lineages based on secretoglobin 1A1 (SCGB1A1) and acetylated tubulin (ACTUB) staining, respectively, as early as 21 DPT and persist to 56 DPT (Fig 1E white and orange arrow heads). Together this indicates that use of the micro-bronchoscope for donor cell delivery does not substantially alter the engraftment or differentiation potential of iBCs in the context of the mouse trachea.

### Intrapulmonary airway delivery of >0.5% PDOC results in significant alveolar hemorrhage and subsequent mortality

Having established that tracheal engraftment of iBCs is not altered by delivery through the MB, we next asked if unilateral intrapulmonary preconditioning was possible using PDOC and the micro-bronchoscope. Delivery of 30 ul 1.75% PDOC into the left mainstem bronchus resulted in 100% mortality within 20 minutes, and gross assessment of freshly explanted lungs revealed severe hemorrhage into the left lobe of 1.75% PDOC treated animals (Fig 2A-B). A dose response of reduced concentrations of PDOC was conducted and survival as a function of time was plotted in a Kaplan-Meier survival curve (Fig 2B).

PDOC concentrations of 1% and lower had an overall survival of greater than 50% at 5-hours (300 minutes) post injury (Fig 2B). Mice that were alive 5-hours post-PDOC injury survived. Histologic analysis was conducted on surviving animals receiving 1% (54.5% survival), 0.5% (94.1% survival), and 0.25% (100% survival) PDOC at 5-hours post-conditioning to assess the degree of alveolar and airway epithelial injury (Fig 2C). At 5-hours post-injury, significant regions of hemorrhage and proteinaceous fluid were noted in the alveolar space of lungs treated with 1% PDOC (Fig 2C2, solid black lines), and to a much lesser degree in lungs treated with 0.5% PDOC (Fig 2C3, solid black lines). Regions of airway epithelial sloughing were present in these lungs (Fig 2C2-3, dashed black lines, asterisks). Minimal alveolar hemorrhage or proteinaceous fluid was noted in the lungs of animals treated with 0.25% PDOC (Fig 2C4, dotted black lines). Contralateral right lobes of injured mice appeared normal (representative image Fig 2C1) as did left lobes of mice receiving PBS (data not shown). We thus conclude that high levels mortality seen in animals treated with concentrations >0.5% is likely a result of acute alveolar damage and alveolar hemorrhage.

### Targeted unilateral delivery of polidocanol through the micro-bronchoscope results in reproducible disrup4on of both airway and alveolar epithelium

The two concentrations of PDOC (0.25% and 0.5%) with >90% survival were selected for further characterization of the airway epithelial injury. At 24-hours post-instillation of 0.25% PDOC minimal airway epithelial cell loss was observed, and the loss was variable between animals (Supp Fig 3). In contrast, mice treated with 0.5% PDOC contained large areas of airway epithelial denudation at both 5- and 24-hours post injury compared to contralateral controls in both the proximal airways and the distal intrapulmonary airways (Supp Fig 4 and Fig 3A). The degree of airway epithelial denudation was quantified at 24-hours post injury and significant denudation was observed compared to contralateral controls (Supp Fig 4B). Interestingly, this persistent airway cell loss at 24-hours post injury is distinct from PDOC-induced injury in the trachea, where basal cell spreading and hyperplasia is seen within 12 hours, leading to complete re-epithelialization by 24 hours and full recovery of a differentiated airway at 7 days post-injury (15, 16). Despite the initial delay in re-epithelialization of the IPAs, the airway epithelial barrier is largely restored by 2 days post injury (Fig 3A,B). At this stage, segments of the airway lacked the normal distribution of mature SCGB1A1+ secretory cells and ACTUB+ ciliated cells, indicating these airways had not yet returned to a pre-injury state. Despite the initial delay in re-epithelialization, the intrapulmonary airway epithelium appears to have fully recovered by 7 days post-injury (Fig 3C).

**Figure 3.**
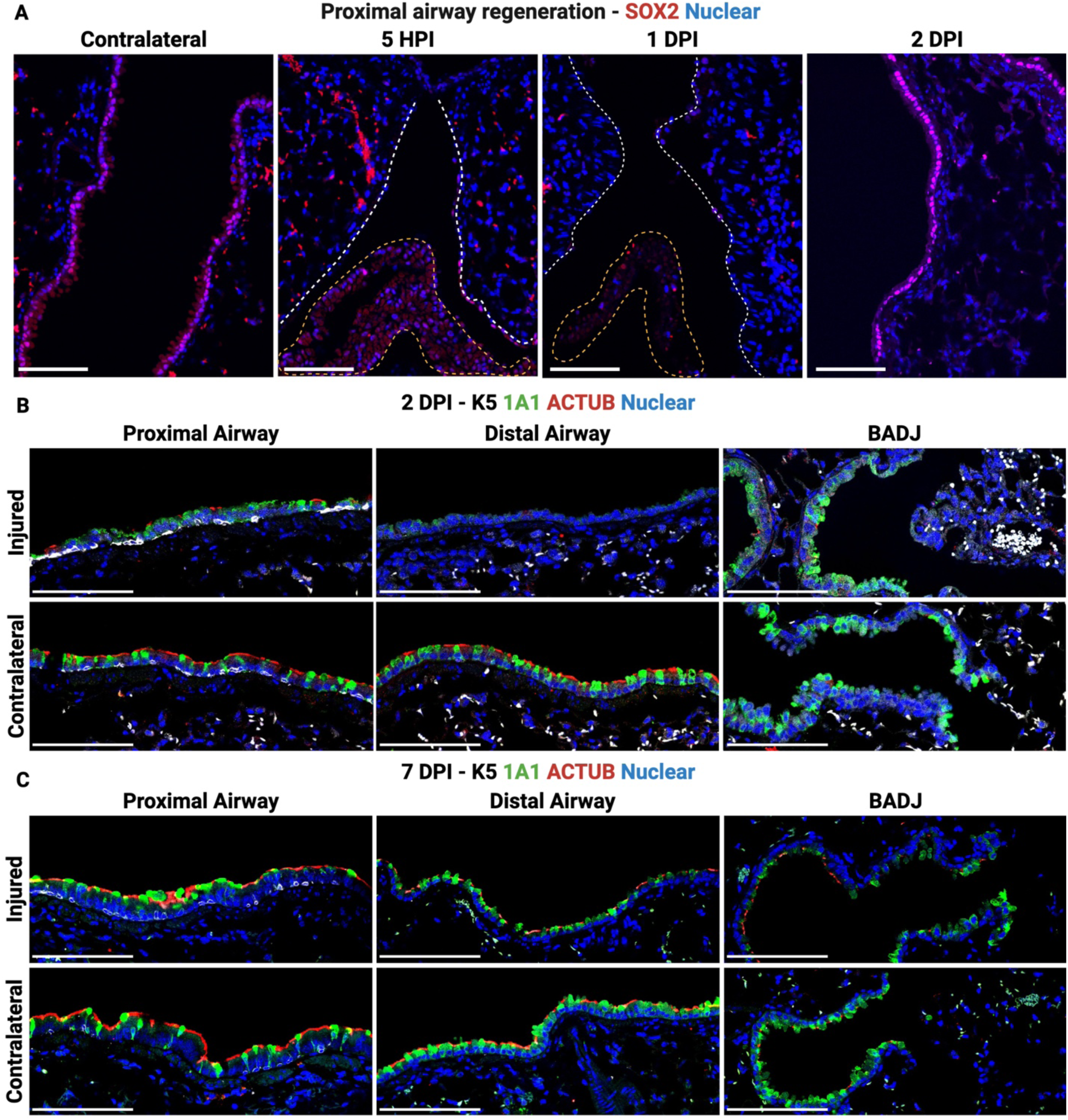
: Recovery of the intrapulmonary airway epithelium following polidocanol injury. (A) Representative immunofluorescence microscopy of intrapulmonary airways at distinct timepoints demonstrating the loss and recovery of airway epithelial cells (SOX2+). White dashed lines indicate denuded airways, and yellow dashed lines indicate sloughed airway epithelial cells. Scale bars are 50 um. (B) Representative immunofluorescence confocal microscopy of intrapulmonary airways at 2 days post-delivery of PDOC with stains for airway lineage markers including K5 (basal cells), ACTUB (ciliated cells), and SCGB1A1 (secretory cells). Scale bars are 100 um. (C) Representative immunofluorescence confocal microscopy of intrapulmonary airways at 7 days post-delivery of PDOC with stains for airway lineage markers including K5 (basal cells), ACTUB (ciliated cells), and SCGB1A1 (secretory cells). Scale bars are 100 um.

**Figure 4.**
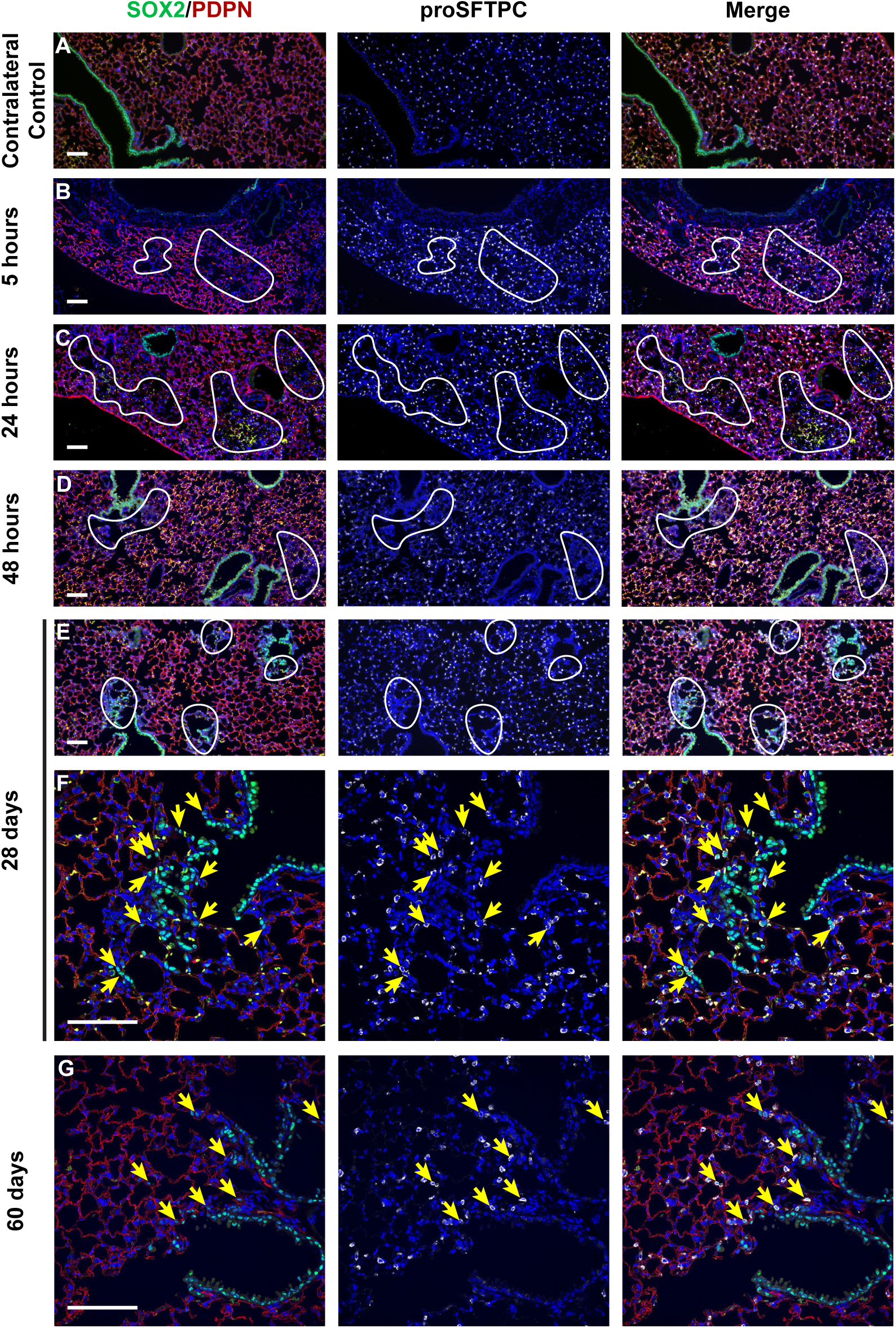
: Alveolar epithelial injury and recovery following injury with 0.5% polidocanol. Representative immunofluorescence microscopy of alveoli and bronchioalveolar duct junctions in lungs following delivery of 0.5% PDOC. Images are taken from a contralateral control (A) or the left lobe at 5 hours (B), 24 hours (C), 48 hours (D), 28 days (E,F) or 60 days (G) after PDOC delivery. White lines outline regions with disrupted alveolar composition as noted by markers of alveolar type II (proSFTPC) and alveolar type I (PDPN) cells. Yellow arrows indicate SOX2+/proSFTPC+ cells. Scale bars are 10tium.

In addition to this airway de-epithelialization, intrapulmonary delivery of 0.5% PDOC induced injury to discrete patches of the alveolar epithelium. As early as 5 hours post injury, proteinaceous debris and alveolar thickening could be found in some alveolar spaces adjacent to bronchioalveolar duct junctions (BADJs) (Fig 2C3). As mentioned, these mice had less extensive injury than in mice treated with 1.0% PDOC (Fig 2C2) and did not demonstrate the substantial hemorrhage seen in mice given higher doses of PDOC (Fig 2A). Analysis of the alveolar injury at 24 hours post-delivery revealed regional but substantial disruption of PDPN+ Alveolar Type 1 (AT1) cells, but not proSFTPC+ Alveolar Type 2 (AT2) cells (Fig 4A-C), especially in regions adjacent to BADJs. However, by 48 hours post injury areas of alveolar thickening were depleted of both AT1 and AT2 cells (Fig 4D). Interestingly, by 28 days post injury these injured areas were repopulated by SOX2+ cells, including a substantial population of alveolar proSFTPC+/SOX2+ cells (Fig 4E,F), which were not seen in uninjured contralateral lungs.. By 60 days post injury alveolar structures had largely returned to normal, but rare alveolar proSFTPC+/SOX2+ cells could still be identified (Fig 4G).

### Primary basal cells are able to engraJ in the intrapulmonary airways following PDOC precondi4oning

Having established that intrapulmonary delivery of 0.5% PDOC results in a non-lethal disruption of the airway epithelium, we next sought to determine whether this was sufficient for subsequent airway epithelial cell transplantation. To generate donor cells congenic to the C57Bl/6J recipient mice, we collected and expanded primary tracheal basal cells from C57Bl/6J mice with a ubiquitous UBC-eGFP reporter, as previously described (15) (Supp Fig 5A,B). Recipient mice were injured in the lei lobe with 3tiul of 0.5% PDOC using the micro-bronchoscopic technique described above. At 24 hours post injury the same methodology was used to deliver 1.0-1.5e6 cells UBC-eGFP primary basal cells as a single cell suspension (Fig 5A and Supp Fig 4). Flow cytometry of an aliquot of cells on day of transplant (DOT) verified that approximately 80% of donor cells were NGFR+;GFP+ basal cells (Supp Fig. 5 D). At 2- and 10- days post transplantation, GFP+ donor-derived cells were clearly detected in the left lobe of recipient mice, but not right lobes based on whole-mount fluorescent microscopy of freshly explanted lungs (Figure 5B).

**Figure 5.**
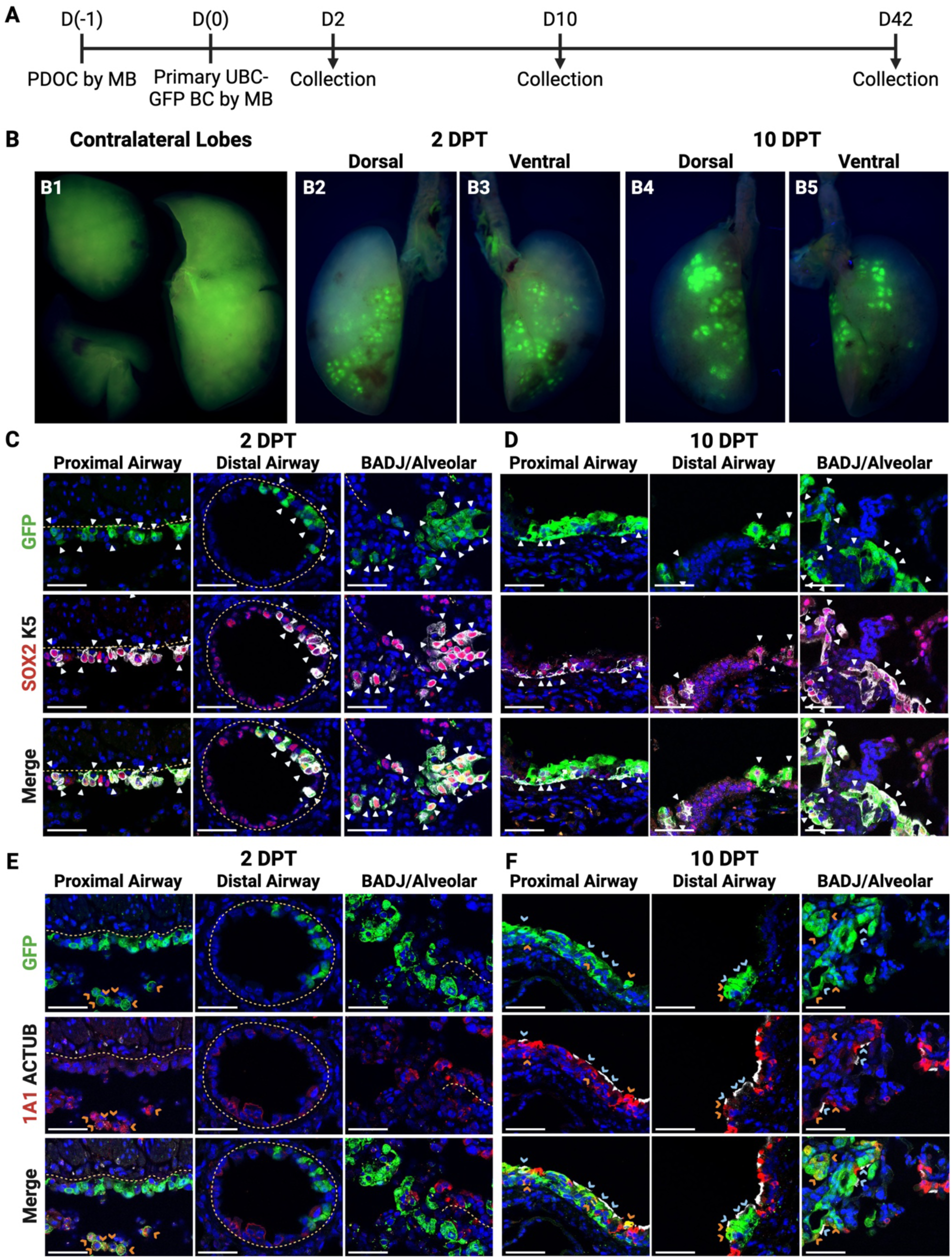
: Intrapulmonary engraJment of primary BCs. (A) Schematic for transplantation of primary GFP+ BCs into mouse lungs. (B) Representative images of non-targeted contralateral lobes (B1) or targeted lobes with GFP+ donor-derived cells at 2 days (B2,B3) or 10 days (B4,B5) post transplantation. (C) Representative immunofluorescence confocal microscopy of donor-derived cells in distinct intrapulmonary regions at 2 days post transplantation, assessing colocalization of GFP with markers of airway epithelium (SOX2) and basal cells (K5, white arrowhead). Scale bars are 50 um. (D) Representative immunofluorescence confocal microscopy of donor-derived cells in distinct intrapulmonary regions at 10 days post transplantation, assessing colocalization of GFP with markers of airway epithelium (SOX2) and basal cells (K5, white arrowheads). Scale bars are 50 um. (E) Representative immunofluorescence confocal microscopy of donor-derived cells in distinct intrapulmonary regions at 2 days post transplantation, assessing colocalization of GFP with markers of secretory cells (SCGB1A1, orange arrowheads) and ciliated cells (ACTUB, blue arrowheads). Scale bars are 50 um. (F) Representative immunofluorescence confocal microscopy of donor-derived cells in distinct intrapulmonary regions at 10 days post transplantation, assessing colocalization of GFP with markers of secretory cells (SCGB1A1, orange arrowheads) and ciliated cells (ACTUB, blue arrowheads). Scale bars are 50 um.

In the OPA tracheal injury and transplantation model described by Ma et al., despite robust engraftment into the trachea, donor-derived cells were only rarely found in the in the intrapulmonary airways, in contrast, we found some level of intrapulmonary engraftment in 100% of transplanted animals. This is despite using a significantly lower number of donor cells per animal (1.-1.5e6 vs 3-6e6 cells) (15). Histologic assessment of engrafted cells focused on three compartments within the transplanted left lobes. First being the proximal intrapulmonary airways, which we define based on the presence of endogenous basal cells and extends distally to approximately the start of the second- generation intrapulmonary airways (32). Second, the distal IPAs, defined as the airways lacking endogenous basal cells; from the second intrapulmonary carina to the bronchoalveolar duct junction (BADJ). Third the alveolar space adjacent to BADJs. GFP+ donor cells were present in all three epithelial compartments, though transplant into the distal IPAs was detected in fewer mice (5/8 mice) with smaller clusters compared to the proximal IPAs (6/7 mice) and alveolar regions (8/8 mice) (Fig. 5C,D,E,F and Supp Fig 6). At two days post-transplant, GFP+ donor-derived cells which had adhered to the basement membrane expressed the pan-airway epithelial cell marker SOX2 and retained expression of the basal cell marker K5 (Fig 5C, white arrowheads). At 2 DPT, when assessed for markers of differentiation to secretory or ciliated lineages, none of the adherent GFP+ donor-derived cells to the basement membrane expressed the secretory marker SCGB1A1 or ciliary marker ACTUB (Fig 5E). GFP+/SCGB1A1+ donor-derived cells could be found within the airway lumen at this timepoint (Fig 5E, orange carrots), however none were detected adhering to the basement membrane. These cells likely correspond to the ∼20% donor cells which were NGFR negative prior to transplantation (Supp Fig 5C), which have previously been identified as secretory-like cells(15).

**Figure 6.**
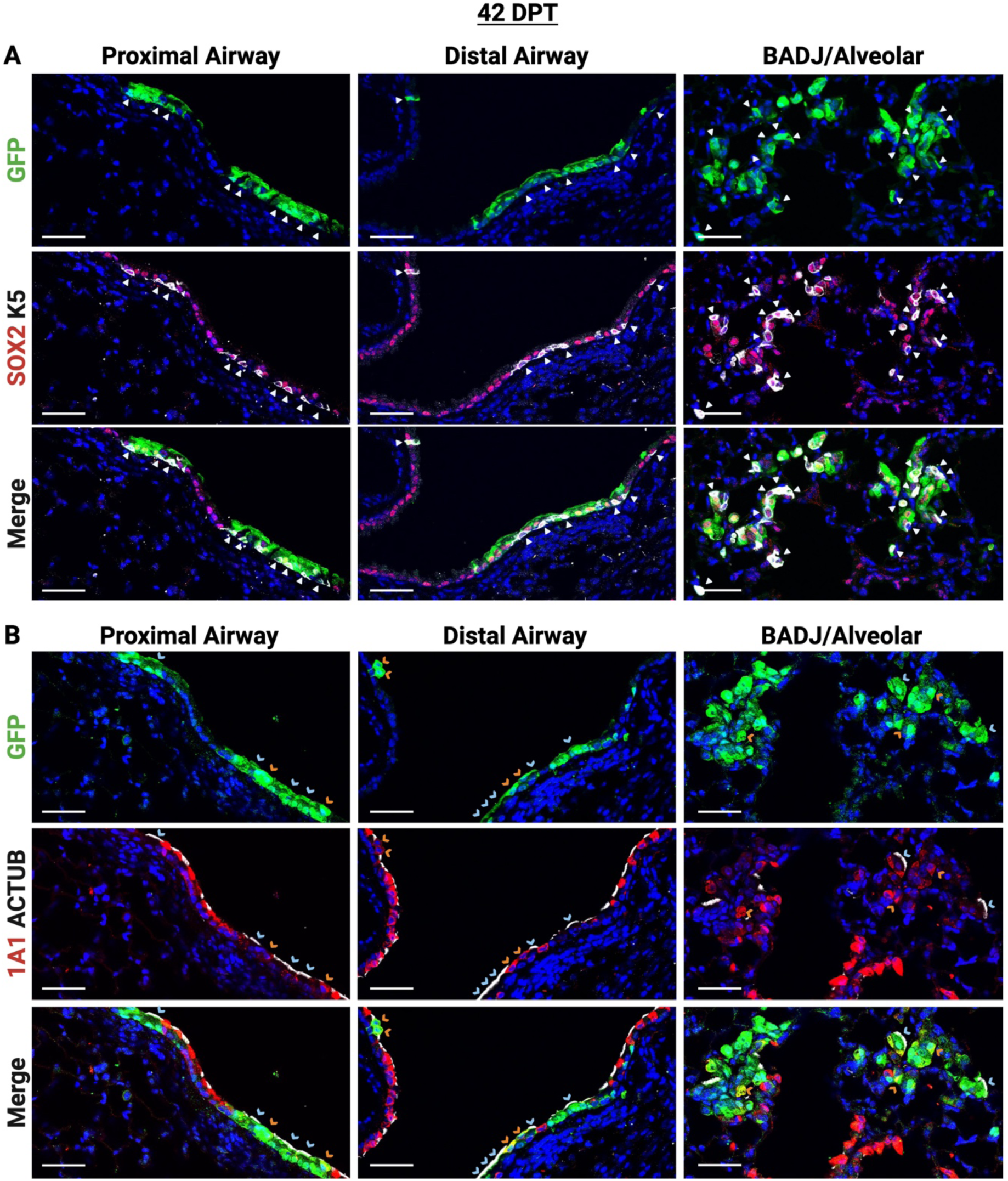
: Persistence of engraJed primary BCs in dis4nct intrapulmonary regions. (A) Representative immunofluorescence confocal microscopy of donor-derived cells in distinct intrapulmonary regions at 42 days post transplantation, assessing colocalization of GFP with markers of airway epithelium (SOX2) and basal cells (K5, white arrowhead). Scale bars are 50 um. (B) Representative immunofluorescence confocal microscopy of donor-derived cells in distinct intrapulmonary regions at 42 days post transplantation, assessing colocalization of GFP with markers of secretory cells (SCGB1A1, orange arrowheads) and ciliated cells (ACTUB, blue arrowheads). Scale bars are 50 um.

At 10 DPT, all GFP+ donor cells maintained expression of SOX2, however, only a subset maintained expression of K5 (Fig 5D, white arrowheads). Interestingly, GFP+/K5+ donor-derived cells are seen in the distal airways as well as the BADJ/Alveolar region where endogenous basal cells are not normally found in mice (Fig 5E, white arrowheads). Furthermore, GFP+ donor-derived cells differentiated into both secretory (SCGB1A1+) and ciliated (ACTUB+) cells in the proximal and distal airways (Fig 5F orange and blue carrot, respectively). GFP+ donor-derived cells in the alveolar region demonstrated similar differentiation into airway lineages (Fig 6F, orange and blue carrots respectively). Even as late as 42 DPT all GFP+ donor-derived cells maintained SOX2 expression with all three regions of engraftment containing cells expressing markers of the three main airway lineages (Fig 6). At all timepoints, we found no proSFTPC (Aleveolar Type 2 cell marker) or RAGE (Alveolar Type 1 cell marker) expression in any donor-derived cells, and thus no evidence of transdifferentiation of GFP+ donor-derived cells in the alveolar region into alveolar cell types (Supp Fig 7).

## Discussion

Cellular engraftment into the lung is in its nascency, and the preconditioning regimens and cells that are sufficient for successful engraftment and amelioration of disease remain major questions. Similar to bone marrow transplantation (33) the bulk of literature has shown that a pre-conditioning or injury step is required to clear endogenous cells from the intended respiratory niche prior to epithelial cell transplantation (15, 18, 21–23, 34–38). These studies have utilized varied methodologies including influenza or *S. pneumonia* infection (37, 39), bleomycin (34–38), acid (0.1N HCl) (37), sulfur dioxide (5), naphthalene (18, 19, 21–23), and detergents including 3-[(3-cholamidopropyl)dimethylammonio]-1- propane sulfonate (CHAPS) (40) and polidocanol (15, 16, 41) to injure one or more disparate sites within the respiratory tract. These pre-conditioning approaches have generally been delivered systemically or broadly within the respiratory tract in order to study lung injury response or as a niche clearing preconditioning step prior to engraftment and have been used without specific regard to the severity of the resultant injury. As a result, these methods, which are useful in a research seong, do not necessarily reflect translational models of cell-transplantation. Intriguingly, a new study has shown engraftment of unfractionated lung cells into the epithelial, endothelial and mesenchymal compartments in a genetic model of idiopathic pulmonary fibrosis without an additional preconditioning step (38). The authors posit that the genetically mediated fibrotic injury created in their model is sufficient to allow donor cells to access the niche. Whether this is borne out in subsequent studies or in other chronic lung diseases remains to be seen.

Here we report a translational micro-bronchoscopic technique for targeted delivery of a preconditioning agent, polidocanol, and subsequent basal cell transplantation to the murine intrapulmonary airways. Through the use of a micro-bronchoscope, we demonstrate precise de- epithelialization of the targeted lobe utilizing the detergent polidocanol with minimal to no impact on the untargeted lobes. With direct instillation into the targeted lobe, a lower concentration of PDOC (0.5% vs 2%) can be utilized resulting in similar airway denudation (Figure 2, 3, S3, and S4) as was seen in in prior studies of tracheal injury (15, 16, 41). Interestingly, we found that intrapulmonary airway denudation by PDOC results in delayed re-epithelialization with bare basement membrane at 24-hours post injury (Figure 4) compared to the tracheal model in which re-epithelialization occurs withing 12 hours (15, 16). The prolonged denudation is a benefit, allowing the PDOC to dissipate and the animals a chance to recuperate from injury prior to undergoing the cell transplantation procedure and likely reflects regenerative differences in the major stem/progenitor populations in the trachea (basal cells) compared to intrapulmonary airways (club cells) (5, 42, 43). Further, we demonstrate that murine primary and PSC-derived basal cells can be delivered through a micro-bronchoscope while retaining their ability to engraft both in the previously reported tracheal model (Figure 1 and S2, (15)) and in our new model of intrapulmonary airway injury (Figs 5 and 6, Supp Figs 5, 6 and 7), and in the case of the intrapulmonary injury model – with higher efficiency than oropharyngeal delivery of cells as described by Ma et al. 2023. Moreover, micro-bronchoscopically delivered basal cells are able to maintain a basal cell pool and differentiate into secretory and ciliated lineages. Interestingly, our studies have shown persistence of donor-derived tracheal KRT5^+^ basal cells in the distal intrapulmonary airways up to 42 DPT (Fig 5, 6, Supp Fig 7). In mice, this region is generally devoid of classical KRT5^+^ p63^+^ basal cells (5, 32), however, multiple groups have described p63^+^ KRT5^nega2ve^ “basaloid” or basal-like progenitor cells residing in this region (39, 44, 45), and in humans’ basal cells are found throughout the extra- and intrapulmonary airways (46). Further, KRT5^+^ p63^+^ cells have been shown to arise in the distal intrapulmonary airways in response to H1N1 influenza infection (44). Therefore, we suggest that the distal intrapulmonary airway is a suitable niche for maintenance of tracheal basal cells at least up to 6- weeks. Whether KRT5^+^ basal cells could persist in this niche long term or appropriately respond to secondary injury remains to be evaluated. Further we find persistence of donor-derived KRT5^+^ primary tracheal basal cells inappropriately within the alveolar space adjacent to the BADJ (Figures 5 and 6) with evidence of differentiation into ciliated and secretory lineages. These findings are congruent with those reported by Ma et al. 2023 with transplantation of ESC-derived basal cells (iBCs) by oropharyngeal aspiration (15). This apparent “bronchiolization” of the alveolar space may be an unintended consequence of the mild alveolar injury created by PDOC. While our mice showed no ill health effects within the 6-weeks of the study, the unintended consequences of the wrong cell in the wrong niche could be devastating, further underscoring the need for robust pre-clinical testing throughout the respiratory tree.

In summary, we have shown that targeted unilateral preconditioning and cell delivery in the murine intrapulmonary airways is possible using the described micro-bronchoscopic technique.

Importantly, this technique will be scalable to larger pre-clinical animal models, where targeted injury and cell delivery could significantly improve the feasibility of cell engraftment experiments by reducing morbidity and delivering the maximum number of cells to the injured location. The persistence and differentiation of airway epithelial cells in the alveolar space adjacent to BADJs highlights the importance of continued robust pre-clinical modeling of cell engraftment in the disparate respiratory compartments and the need to better understand the factors governing cell-niche interactions. Ultimately, these models will help advance our knowledge of intrapulmonary airway and alveolar repair and engraftment which may lead the way to cell-therapies for genetic airway diseases.

## Acknowledgements

We thank Brian R. Tilton, Anna C. Belkina MD, PhD and the entire BUSM Flow Cytometry Core, as well as Hans Gertje and Nicholas Crossland, DVM of the BUSM Integrated Biomedical Imaging Service (NIH S1TIOD030269). We would like to thank the members of the Boston University Pulmonary Center for their support and suggestions throughout the course of this research. Additionally, we are indebted to our colleagues at the Center for Regenerative Medicine, especially Gabrielle Cockerham and Meenakshi Lakshminarayanan for administrative support, Mary Lou Beermann, Greg Miller, PhD, and Amelia Jay for overall laboratory support, and Marianne James for reprogramming and characterization of iPSC lines.

## Author contribu4ons

Project conceptualization, A.K. and F.J.H.; mouse donor cell generation, A.K. and L.M.; mouse injury and cell transplantation A.K., M.J.H., L.M., B.R.T., and P.B.; injury and transplantation recipient analysis A.K., M.J.H., and P.B., writing A.K., M.J.H. and F.J.H., supervision D.N.K. and F.J.H.

---

**Supplemental Figure 1.**
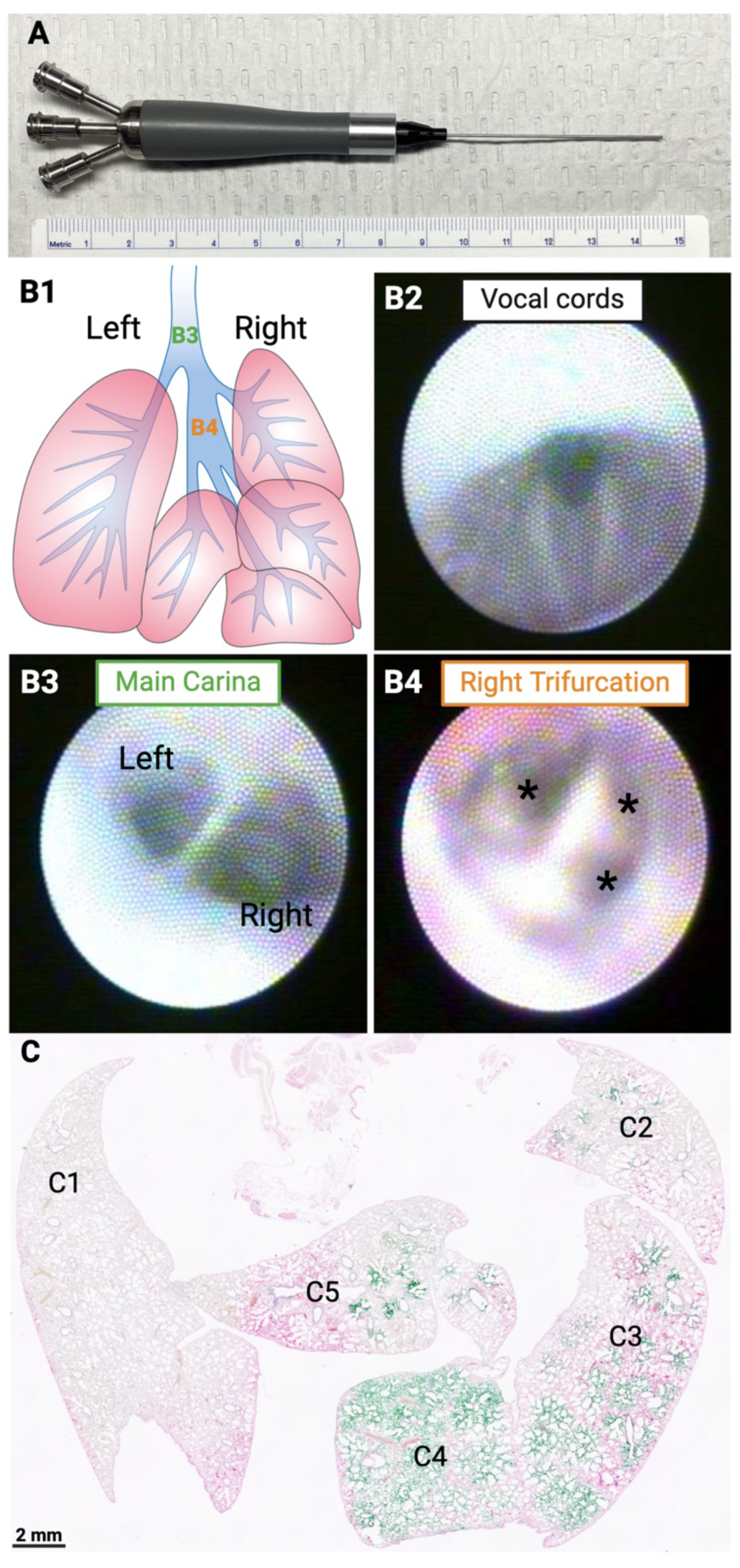
: Unilateral fluid delivery through micro-bronchoscopy (A) Image of the handle and sheath of the micro-bronchoscope used in these studies. (B) Schematic of the mouse lung (B1) along with images taken with the micro-bronchoscope at the vocal cords (B2), main carina (B3), and right trifurcation (B4). (C) Representative H&E stain of lung lobes following delivery of of green dye into the right lobes.

**Supplemental Figure 2.**
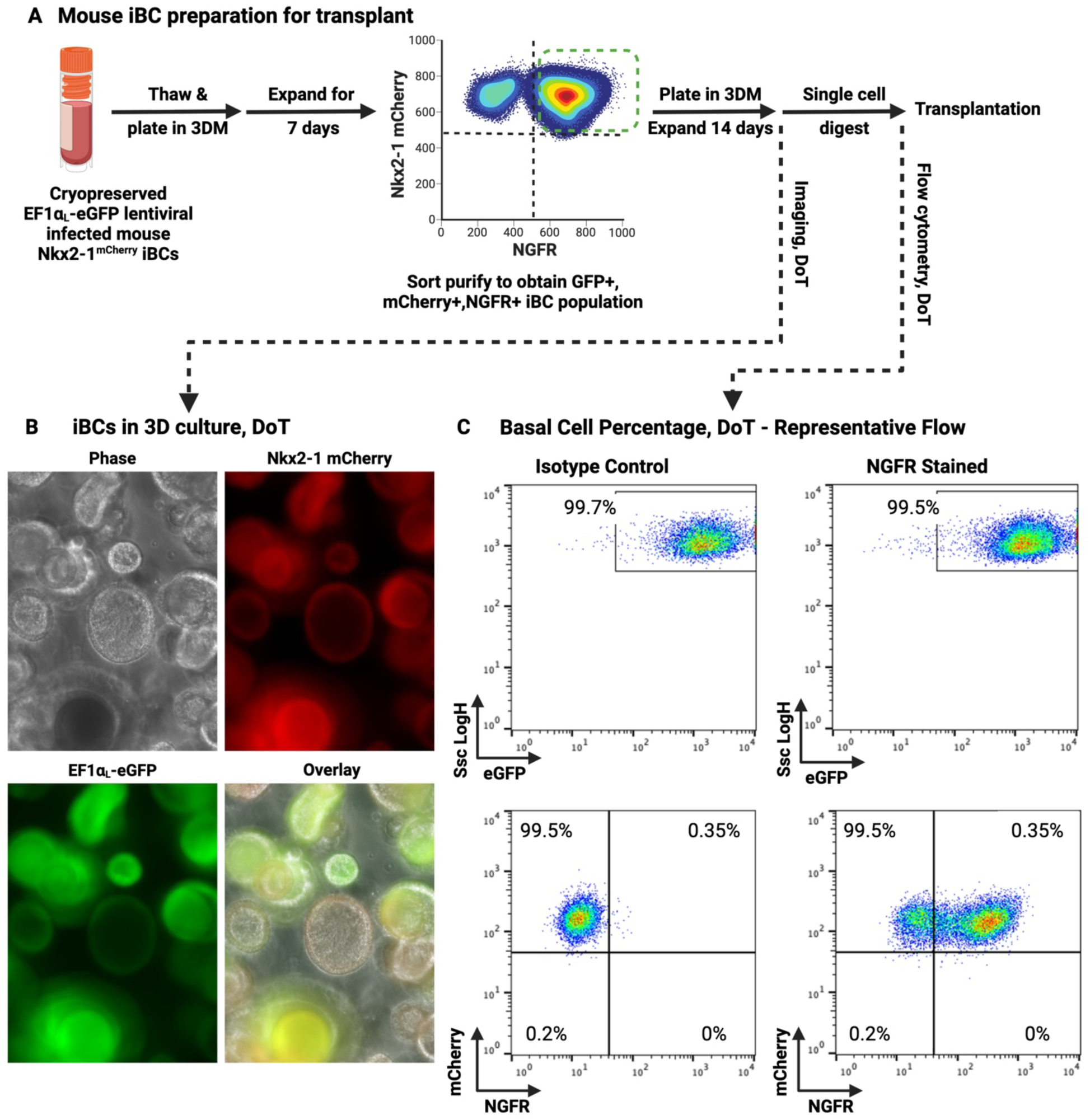
: Culturing and preparing donor mouse iBCs. (A) Schematic of the culturing, FACS purification, and digestion of mouse iBCs. (B) Representative images of cells grown for 14 days following FACS purification. (C) Representative flow assessment of cells grown for 14 days following FACS purification with staining for either NGFR or an isotype control.

**Supplemental Figure 3.**
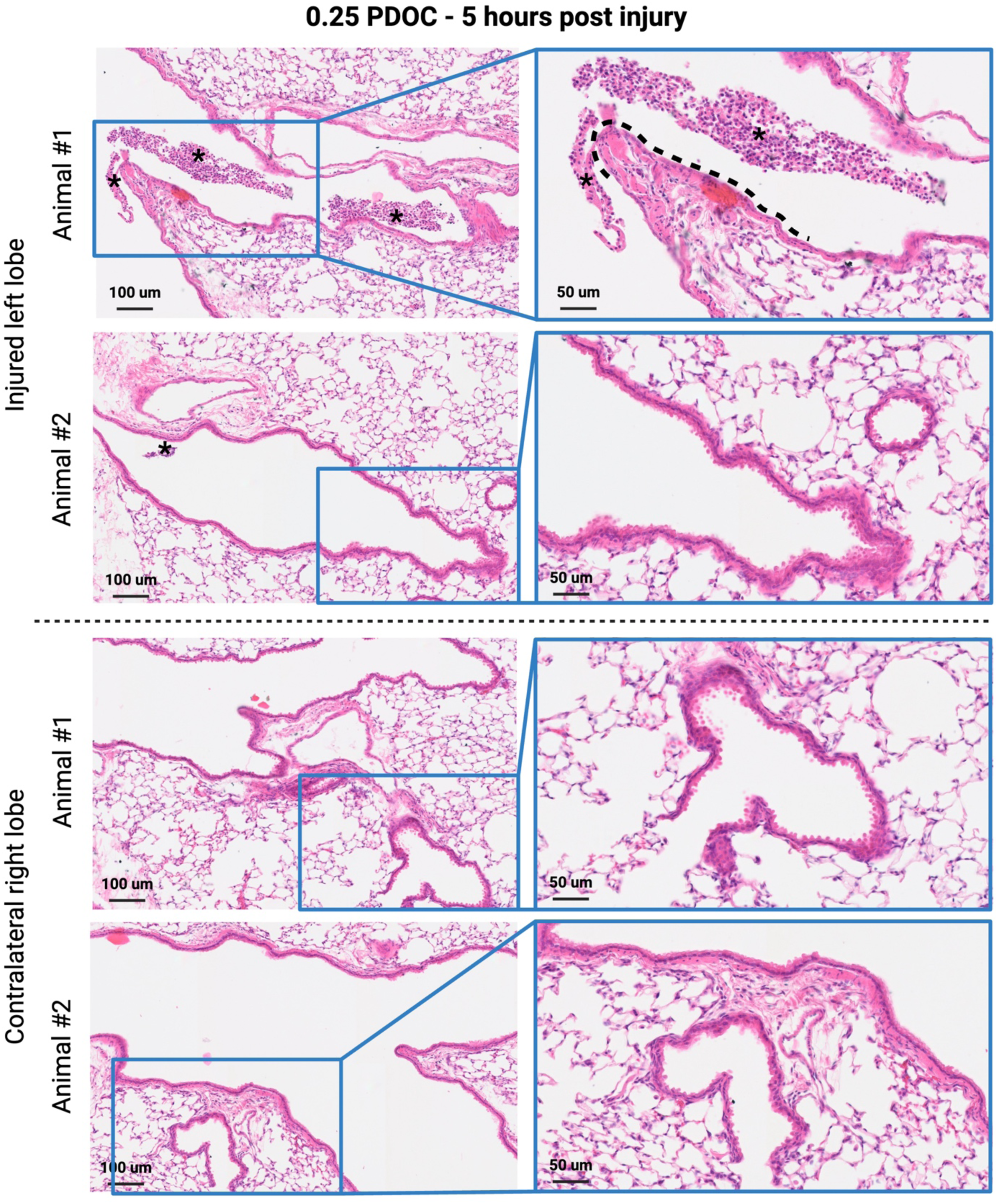
: Intrapulmonary airway epithelial damage at 24 hours post 0.25% polidocanol Representative H&E images of the injured left lobe and contralateral control lobe at 24 hours post-delivery of 0.25% PDOC highlighting minimal denuded airway epithelium (dashed line) and sloughed cells (asterisk).

**Supplemental Figure 4.**
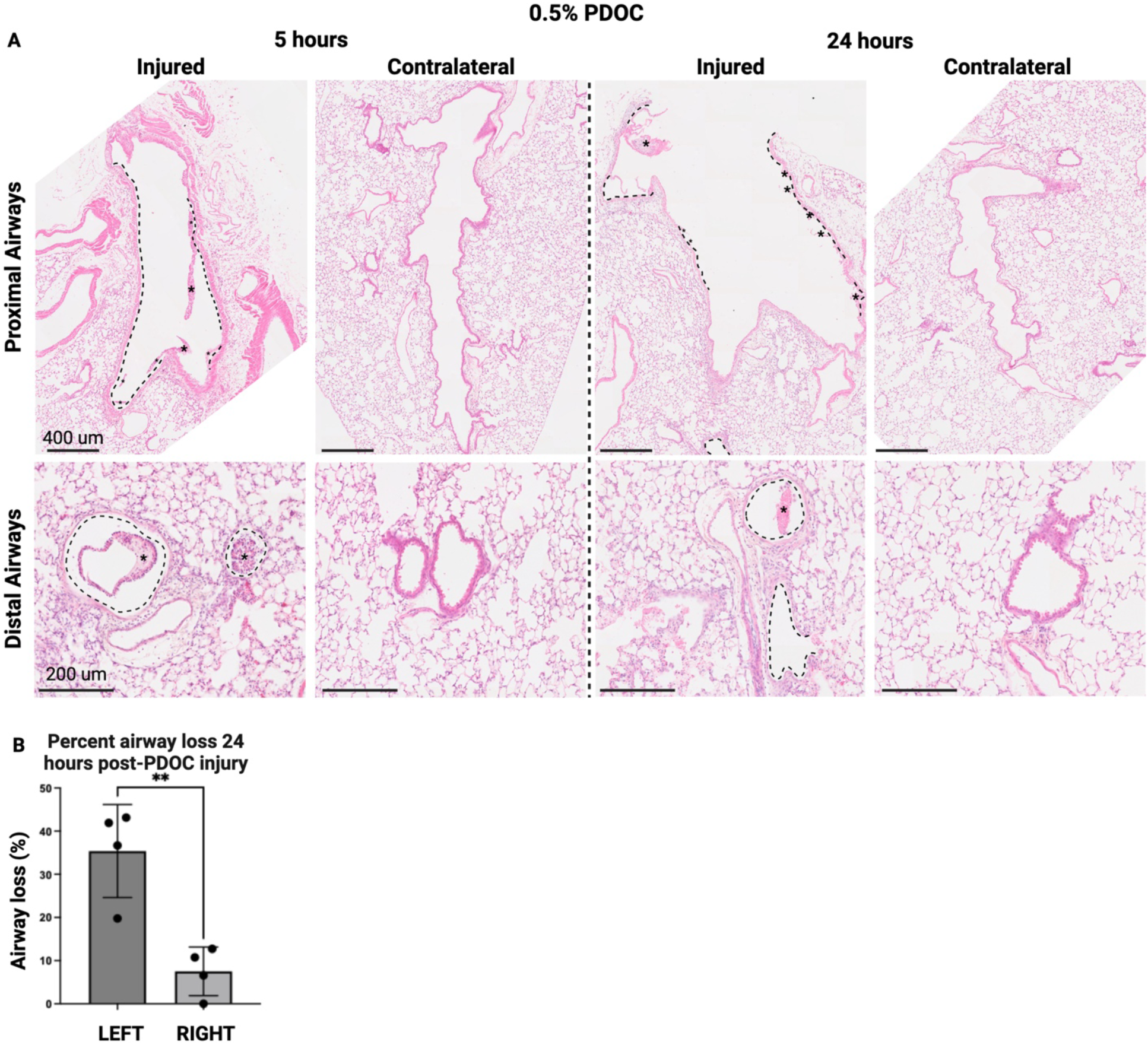
: Intrapulmonary airway epithelial damage at 5 and 24 hours post 0.5% polidocanol (A) Representative H&E images of the injured left lobe and contralateral control lobe at 5 and 24 hours post-delivery of 0.5% PDOC highlighting denuded airway epithelium (dashed line) and sloughed cells (asterisk). (B) Quantification of denuded airways in left and right lobes at 24 hours post-delivery of 0.5% PDOC. n=4 mice per condition.

**Supplemental Figure 5.**
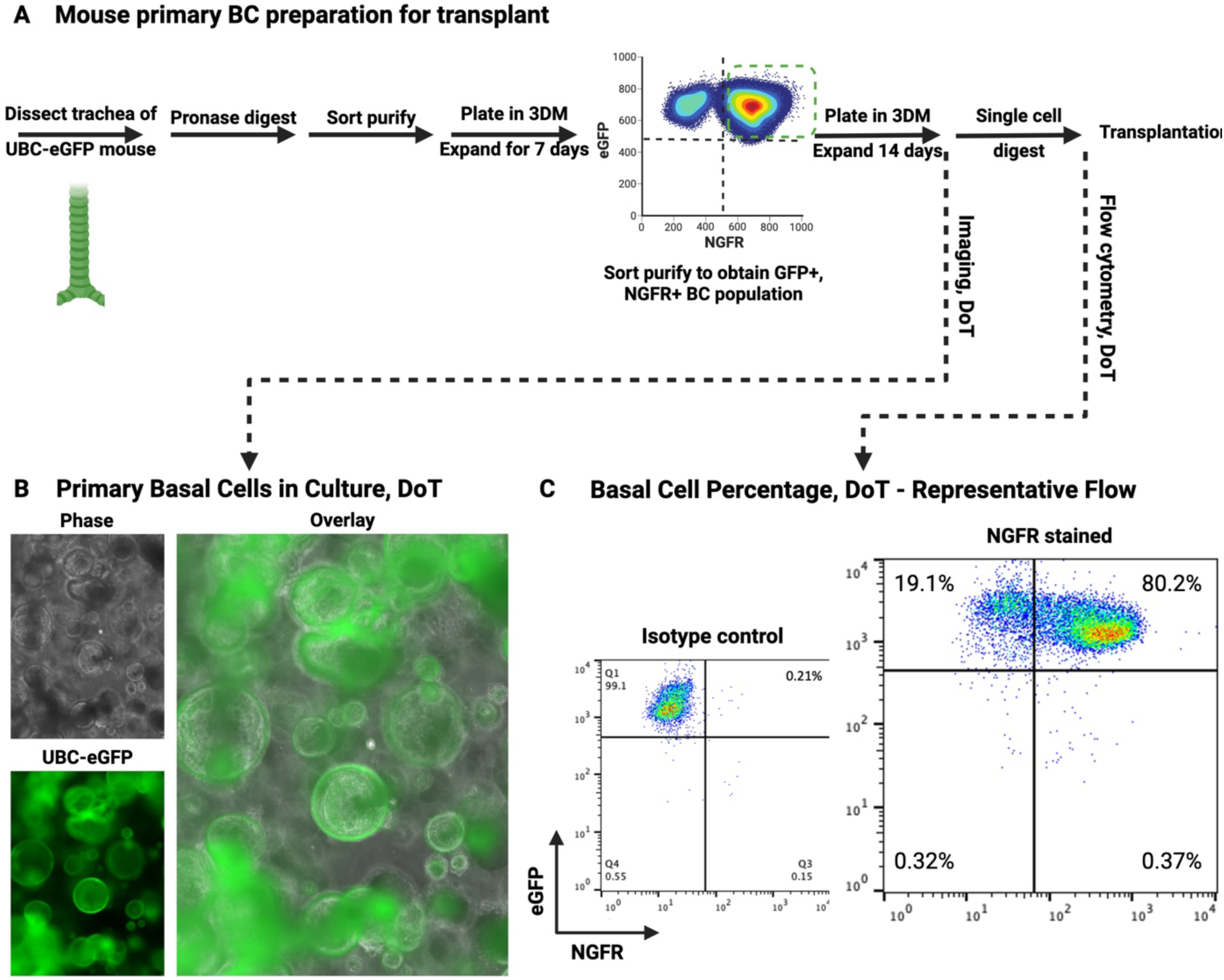
: Culturing and preparing donor mouse primary BCs. (A) Schematic of the collection, culturing, FACS purification, and digestion of mouse primary BCs. (B) Representative images of cells grown for 14 days following FACS purification. (C) Representative flow assessment of cells grown for 14 days following FACS purification with staining for either NGFR or an isotype control.

**Supplemental Figure 6.**
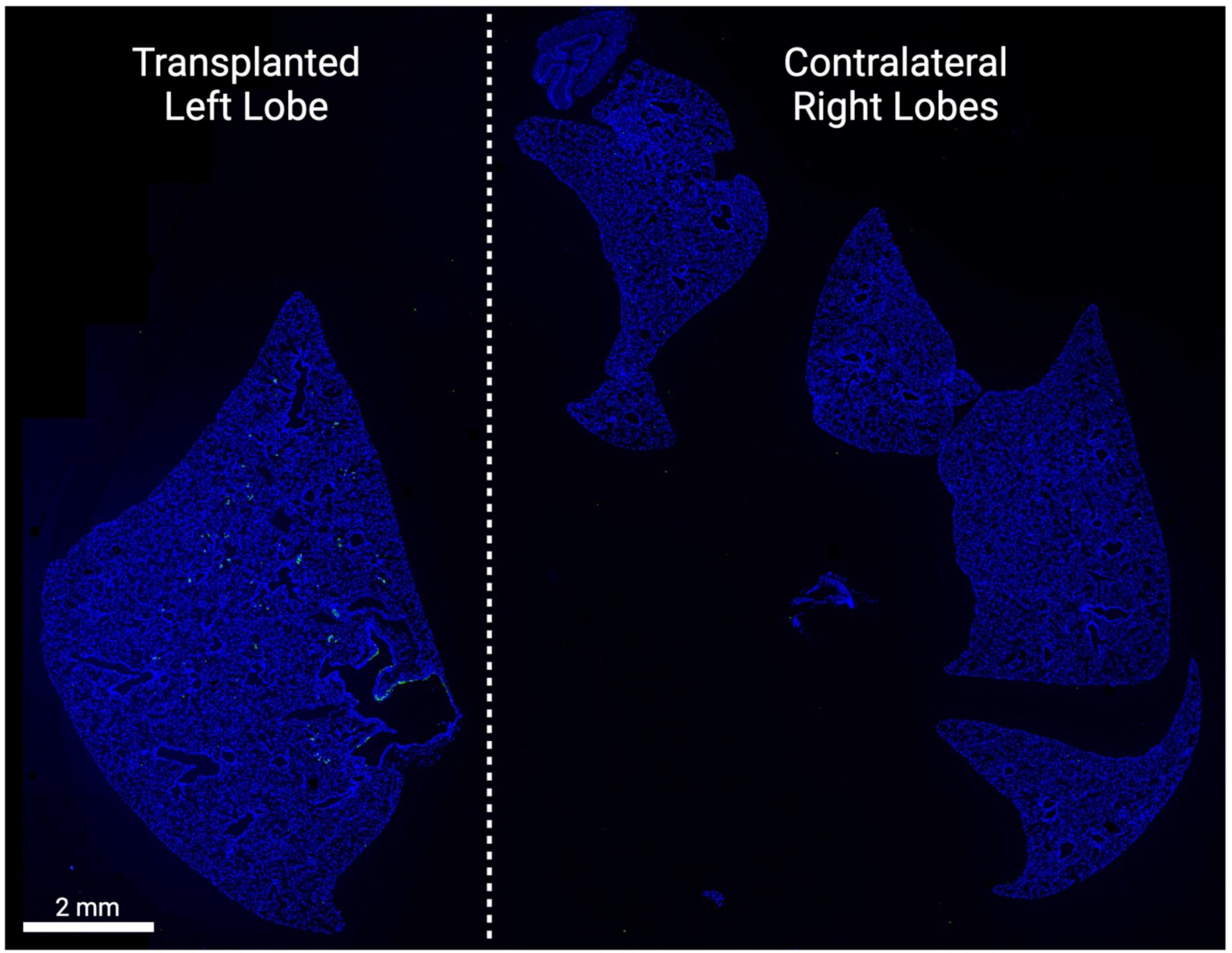
: Low magnifica4on image of intrapulmonary engraJment following targeted polidocanol precondi4oning Representative immunofluorescence microscopy of lung lobes at 56 days post engraftment of GFP+ primary BCs.

**Supplemental Figure 7.**
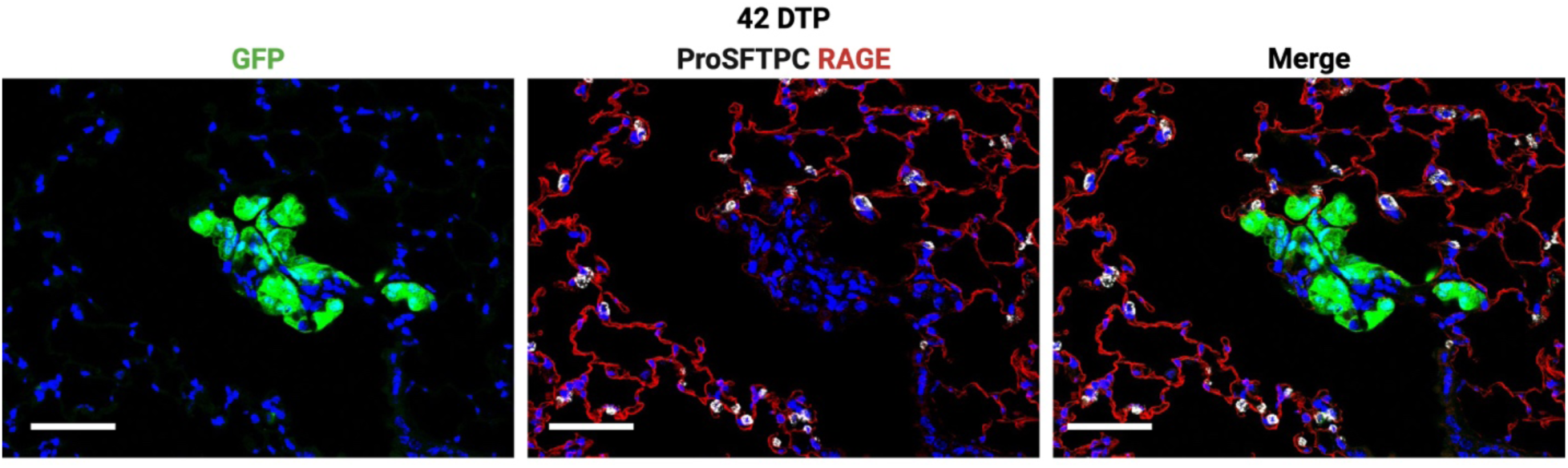
: EngraJed primary BCs do not give rise to alveolar lineages Representative immunofluorescence confocal microscopy of alveolar donor-derived cells at 42 days post transplantation, highlighting that GFP+ donor-derived cells do not express markers of alveolar type II (proSFTPC) or alveolar type I (RAGE) cells. Scale bars are 50 um.

**Supplemental Video 1:** Visualiza4on of major airways through micro-bronchoscopy Representative video taken with the micro-bronchoscope and identifying major airways of the mouse lung.

**Supplemental Table 1:**
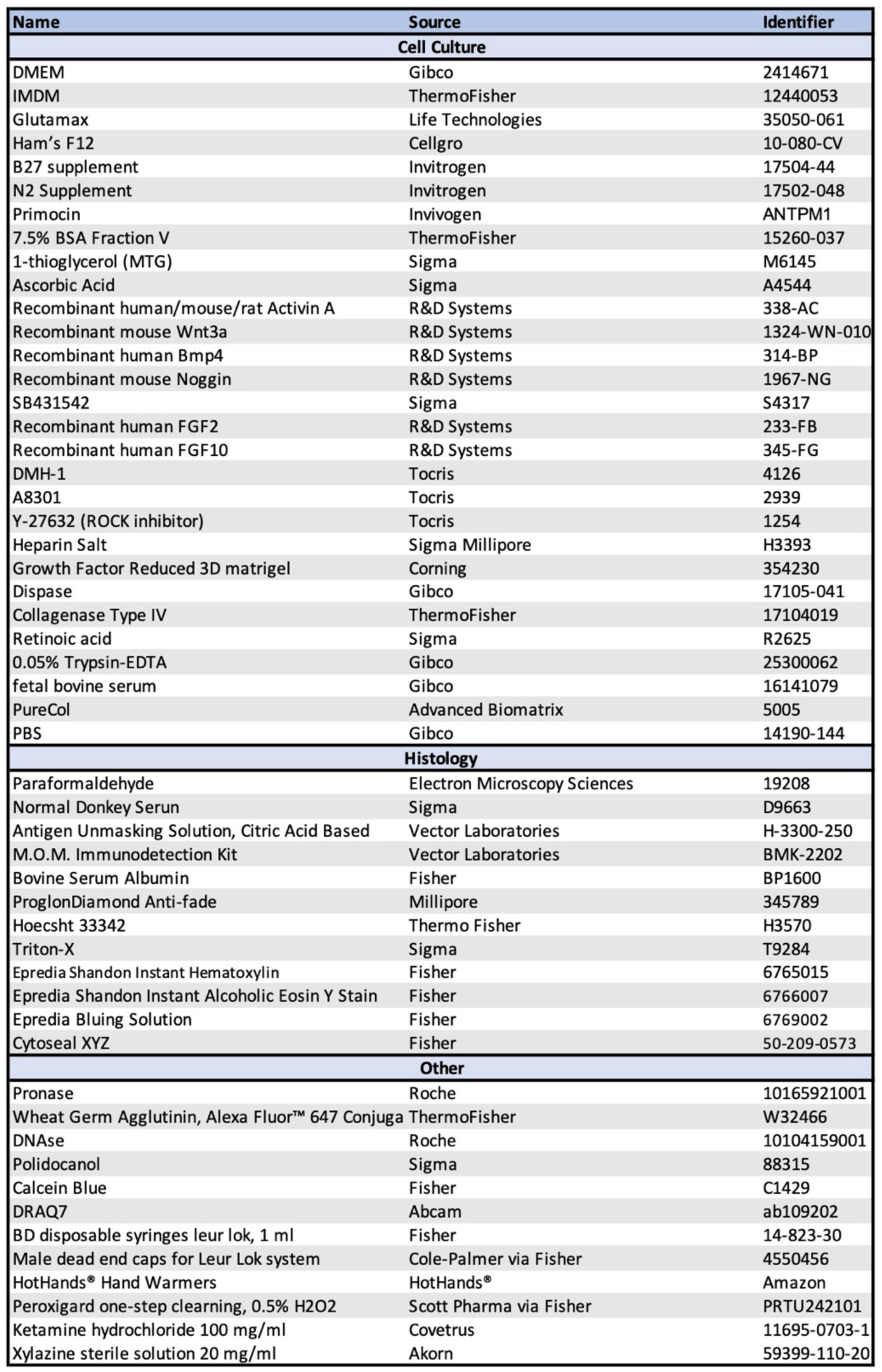
Reagents

**Supplemental Table 2.** : An4bodies

| **Name** | **Dilution** | **Source** | **Identifier** |
| --- | --- | --- | --- |
| Mouse Anti-NGFR Antibody | 1:1000 | Abcam | ab245134 |
| BV421 Rat Anti-Mouse CD326 | 1:500 | BD Biosciences | 563214 |
| Goat Anti-GFP Antibody | 1:500 | US Biological | G8965-01E |
| Chicken Anti-GFP Antibody | 1:200 | Aves Labs | GFP-1020 |
| Rat Anti-GFP Antibody | 1:500 | Abcam | ab252881 |
| Rabbit Anti-NKX2-1 Antibody | 1:200 | Abcam | ab76013 |
| Rabbit Anti-KRT5 Antibody | 1:500 | BioLegend | 905501 |
| Mouse Anti-Acetylated-Tubulin Antibody | 1:1000 | Sigma | T7451 |
| Goat Anti-SCGB1A1 (human) Antibody | 1:1000 | Sigma | ABS1673 |
| Rat Anti-SOX2 Antibody | 1:100 | Invitrogen | 14-9811-82 |
| Rabbit Anti-SFTPC Antibody | 1:500 | Abcam | ab211326 |
| Hamster Anti-PDPN Antibody | 1:200 | Invitrogen | 14-5381-82 |
| Goat Anti-RAGE Antibody | 1:100 | R&D systems | AF1145 |
| Alexa Fluor 488 donkey anti-chicken IgY (IgG) (H+L) | 1:500 | Jackson Immuno Research | 703-545-155 |
| Alexa Fluor 488 donkey anti-goat IgG (H+L) | 1:500 | Jackson Immuno Research | 705-545-003 |
| Alexa Fluor 488 donkey anti-rat IgG (H+L) | 1:500 | Jackson Immuno Research | 712-546-153 |
| Alexa Fluor 488 donkey anti-rabbit IgG (H+L) | 1:500 | Jackson Immuno Research | 711-546-152 |
| Alexa Fluor 488 donkey anti-mouse IgG (H+L) | 1:500 | Jackson Immuno Research | 705-545-150 |
| Alexa Fluor 546 donkey anti-Rabbit IgG (H+L) | 1:500 | Thermo Fisher | A11055 |
| Alexa Fluor 568 donkey anti-mouse IgG (H+L) | 1:500 | Thermo Fisher | A10037 |
| Cy3 donkey anti-rat IgG (H+L) | 1:500 | Jackson Immuno Research | 712-166-153 |
| Cy3 donkey anti-rabbit IgG (H+L) | 1:500 | Jackson Immuno Research | 711-165-152 |
| Cy3 donkey anti-goat IgG (H+L) | 1:500 | Jackson Immuno Research | 705-165-147 |
| Alexa Fluor 647 donkey anti-rabbit IgG (H+L) | 1:500 | Thermo Fisher | A32795 |
| Alexa Fluor 647 donkey anti-goat IgG (H+L) | 1:500 | Jackson Immuno Research | 705-605-003 |

